# Biofeedback speeds adaptation to exoskeleton gait assistance

**DOI:** 10.1101/2025.06.21.660857

**Authors:** Ava Lakmazaheri, Steven H. Collins

## Abstract

Exoskeletons may enhance mobility, but users require extensive training to receive their full benefit. While augmented feedback can accelerate motor learning, its application to exoskeleton-assisted gait is limited by the complexity of locomotor function and human-robot interaction. We developed a visual biofeedback system to guide novice users of an ankle exoskeleton to modify their ankle joint kinematics and foot placement toward patterns associated with improved energy economy. Biofeedback-based training doubled the energy savings from exoskeleton use, enabling people to achieve benefits comparable to fully adapted users in one-quarter of the time. Notably, participants in this study had not fully adapted after the hour-long training session, underscoring the task’s difficulty and suggesting that greater benefit from exoskeletons may be unlocked with continued use of this approach. Energy savings were associated with increased exploration and progression toward lower-cost gait parameters in task-relevant dimensions. Our findings demonstrate that biofeedback can accelerate motor adaptation to exoskeletons, potentially enhancing their effectiveness and promoting broader device adoption.

## Introduction

Lower-limb exoskeletons have emerged as a promising tool to enhance gait for people experiencing mobility decline. By reducing the energy cost of walking, exoskeletons may help individuals sustain movement for longer periods without added exertion. This greater gait capacity can promote community engagement and physical activity, resulting in higher quality of life and increased longevity [1,2].

Slow motor adaptation is a limiting factor in the study and application of exoskeletons. Biomechanical and physiological outcomes during exoskeleton use change substantially over the course of motor adaptation [3–8]. Fully adapted gait behaviors, emerging after several hours of use [3–5], are rarely reflected in laboratory-based studies. In real-world contexts, reduced effectiveness during early experience with an exoskeleton may decrease people’s expectancies of the device, which can negatively impact both motor performance and the likelihood of longer-term use [9,10].

Long adaptation times reflect the complexity of underlying motor processes. Adaptation involves the exploration of candidate coordination patterns preceding discovery and exploration of an optimal solution [4,11–13]. As the body is an over-actuated system with control over thousands of motor units, this exploration occurs in a high-dimensional space. To explore more efficiently, the nervous system attempts to identify task-relevant dimensions before operating within a refined search space [4,14].

Use cases where exoskeletons have the greatest potential impact may involve even longer timescales of motor adaptation. Age-related or neurological gait impairments can increase motor noise, impeding the evaluation of coordination patterns necessary to reduce search space dimensionality [15–17]. Additionally, real-world exoskeleton use requires parsing task-relevant information from an increased number of sensory inputs while greater demands are placed on neural resources [13]. As exoskeleton research progresses to these critical use cases, it is essential to identify reliable methods of accelerating motor adaptation.

Skills may be acquired more quickly through well-structured practice and provision of augmented feedback. Training experiences that induce moderate variability in goal or task execution can speed learning by highlighting variables that link actions to goal behaviors [12,14,18]. This training approach has been found to facilitate faster learning than self-guided exploration [19]. Augmented feedback can also support learning by highlighting differences between actual and desired movements and the direction of required behavioral change [13]. Many modalities and delivery schedules of biofeedback have been explored; its effectiveness can vary largely with factors related to both the task and the learner [20,21].

Structuring practice is challenging for locomotor skills, especially in the context of assistive device use. Motor learning research has predominantly focused on well-defined tasks such as reaching, where parameters like response time, task difficulty, and performance are more easily quantified and experimentally controlled. In contrast, the rapid, cyclic nature of gait provides a narrow, fixed window over which an individual must process feedback, execute changes, and evaluate their impact before the next step begins. While exoskeletons are often designed to improve specific performance outcomes such as energy consumption, there is no standardized execution of movement that reliably achieves this goal. Performance outcomes for gait also require many steps to evaluate reliably, in contrast to typical outcomes for upper-limb movements that can be assessed continuously or on a per-reach basis. Characterizing task dynamics are further complicated by our limited understanding of physical human-exoskeleton interaction. Altogether, these factors impede our ability to develop a formal system that can guide individuals toward desired gait behaviors.

To address some of these challenges, we previously characterized gait changes as novice exoskeleton users discovered their energy-optimal gaits while walking with plantarflexion assistance [5]. While people are immediately able to take advantage of exoskeletons by offloading assisted muscle activity and increasing ankle power during push-off, they also experience gait perturbations that were less energy-efficient. Novices responded to exoskeletons by pre-emptively plantarflexing as exoskeleton torque rose early-to-mid-stride and changed foot placement strategies, likely to increase stability. Motor adaptation over multiple hours corresponded to these gait characteristics returning toward values observed during typical, unperturbed gait. It is these multi-hour changes that are associated with users tripling their metabolic savings with exoskeletons [3,4].

Visual biofeedback has been used to modify related gait behaviors in the absence of a robotic device. Studies have induced changes in spatiotemporal gait parameters, muscle activation, and joint kinematics using feedback with varying levels of information provided, from abstract illustrations to displays of real-time trajectories every gait cycle (see [22–26] for review and examples). Fewer studies have explored the use of biofeedback in conjunction with lower-limb exoskeletons. In some cases, it has been used as a complementary intervention: biofeedback with a resistive exoskeleton has led to greater changes in step length and plantarflexor activity than with an exoskeleton alone [27,28]. Biofeedback designed to directly influence exoskeleton interaction has been less common; recent studies have demonstrated that visual cues could alter total ankle torque [29] and reduce soleus activity [30] with an exoskeleton assisting plantarflexion. It may be helpful to additionally target gait kinematics and metabolic cost to enhance adaptation.

In this study, we investigated the effect of biofeedback-based training on the rate and extent of learning to walk with an assistive ankle exoskeleton. Our approach guided novice exoskeleton users toward biomechanical gait changes previously determined to be relevant for the task of energy optimization [5]. We assessed metabolic and kinematic changes compared to matched controls who underwent exoskeleton training without augmented feedback. We expected that training with biofeedback would promote gait exploration toward desired behaviors, corresponding to greater energy savings from assistance within one hour of exposure.

## Methods

### Participants

Twenty-six healthy young adults naïve to exoskeleton use participated in this study. Thirteen participated in exoskeleton training with custom visual biofeedback (5 F, age: 23 ± 4 years, height: 1.76 ± 0.10 m, mass: 69.6 ± 11.9 kg). Thirteen demographically matched controls underwent the same duration of training without biofeedback (5 F, age: 24 ± 5 years, height: 1.74 ± 0.10 m, mass: 67.7 ± 10.9 kg). The study protocol was approved by the Stanford University Institutional Review Board and all participants provided written informed consent before participation.

### Experimental Setup

Participants walked on an instrumented treadmill at 1.25 m/s while wearing a respirometry mask and bilateral ankle exoskeletons providing plantarflexion assistance (Fig. 1A). Lightweight, tethered exoskeletons [31] applied a single-peak plantarflexion torque once per step, parameterized by a peak magnitude of 0.54 times body mass at 52.9% stride with a rise time of 26.2% and a fall time of 9.8% stride. This profile was previously found widely beneficial for energy economy [3,32] and was used in prior characterization of expert exoskeleton use [5]. Ankle angle was measured at 1000 Hz from a rotary encoder (Renishaw) mounted at the exoskeleton ankle joint. Step length and width were calculated during walking from the center of pressure at foot contact via treadmill force plate data (Bertec). Metabolic rate was calculated using indirect calorimetry (Cosmed Quark CPET) and the Brockway equation [33]. Energy cost was computed as the difference between instantaneous metabolic rate during walking and resting metabolic rate during quiet standing, normalized by body mass. Participants were instructed to fast for four hours before the start of the session to mitigate the thermic effect of food on respiratory data.

**Figure 1.**
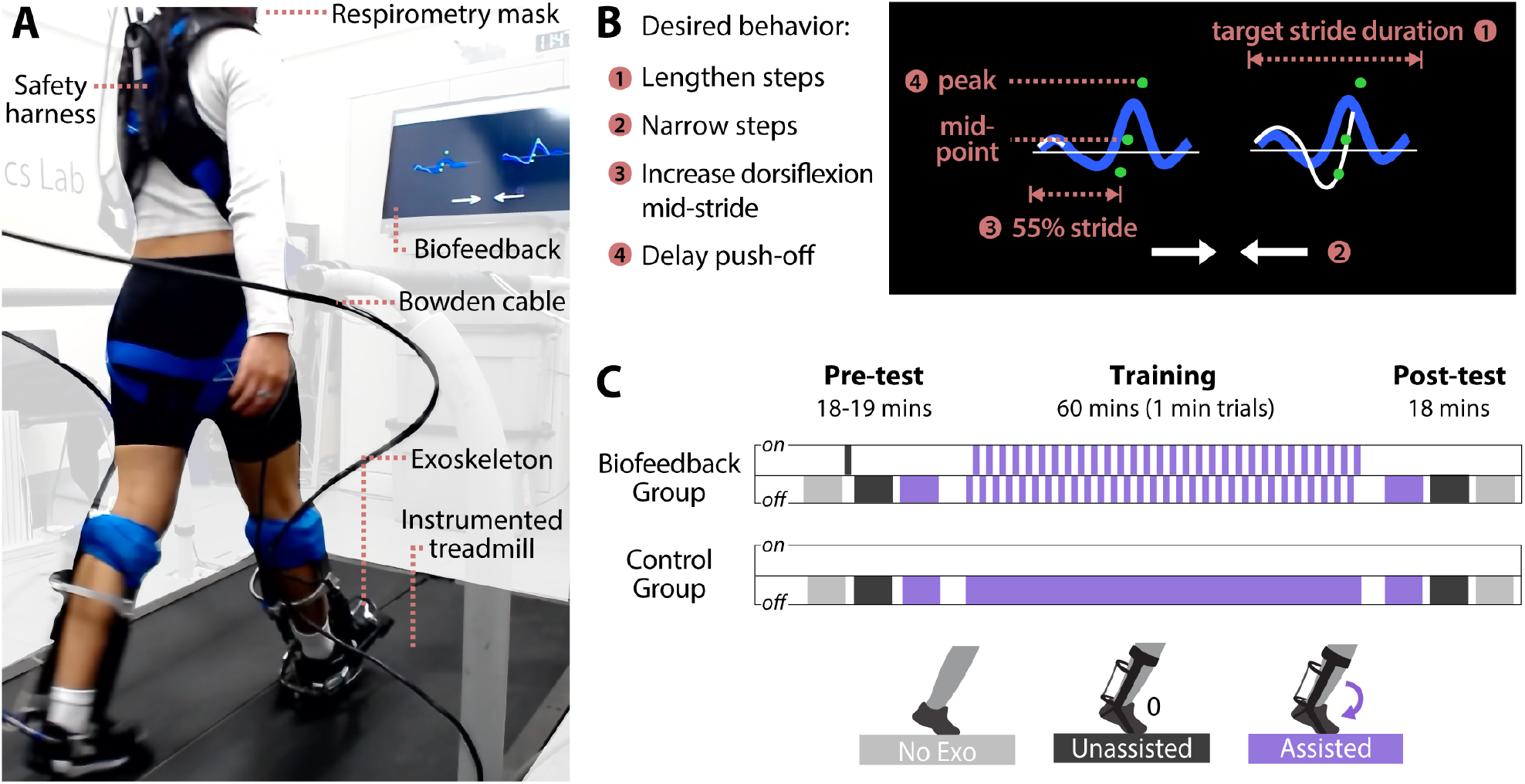
Experimental setup and protocol. **(A)** Photograph of experimental setup. Participant wears tethered bilateral ankle exoskeletons and a respirometry mask (not pictured) while walking on an instrumented treadmill in front of a screen displaying biofeedback. **(B)** Illustration of visual biofeedback. Average (blue) and real-time (white) ankle angle were plotted. Participants were instructed to pass curves through three target nodes (green), corresponding to desired changes in step length and ankle joint kinematics. Inward-pointing arrows were displayed at the bottom of the screen when steps exceeded the desired width. **(C)** Schematic of experimental protocol. Participants completed a pre-test, 60 minutes of group-specific training (with or without biofeedback), and a post-test to evaluate training effects.

A visual biofeedback system was designed to simultaneously increase step length, decrease step width, increase mid-stride dorsiflexion angle, and delay the timing of peak plantarflexion (Fig. 1B). Data were processed from a real-time computer (Speedgoat) and displayed in a custom MATLAB figure. Real-time ankle angle and an average profile of the last 10 strides were plotted bilaterally (angular resolution: 0.001 rads, temporal resolution: 0.001 sec). Left and right plots were cleared on respective heel strikes. Three target nodes were displayed, corresponding to desired (1) ankle angle at 55% stride, (2) midpoint between minimum and maximum angle, and (3) peak plantarflexion angle. The region spanning 50-65% of stride changes broadly as people adapt to exoskeletons; we selected these three points given feedback about visual clarity during pilot testing and the consistency of novice versus expert trends for these features in prior work [5]. Nodes were scaled as a percent of desired stride duration to prompt corresponding changes in step length. To cue a narrower stance, inward-pointing arrows were displayed when current step width exceeded the desired value.

### Experimental Protocol

A pre-test was conducted with six-minute trials of standing at rest, walking with no exoskeletons, walking with exoskeletons providing zero torque, and with exoskeletons providing static torque (Fig. 1C). Trials were ordered to minimize exoskeleton don/doff time. Both groups were given the instruction to “walk comfortably and let the device do the work for you” before the first assisted trial. The control group was given no further instructions on how to walk with exoskeletons.

The biofeedback group underwent a short demonstration period after first donning the exoskeletons. Participants were encouraged to explore gait changes and observe how these changed the display, with the goal of passing plotted curves through the green nodes and clearing the arrows from the screen. Participants practiced the task while wearing exoskeletons providing zero torque for approximately one minute and had the opportunity to ask clarifying questions about the display.

Each group then underwent 60 minutes of training with static assistance. Short breaks were provided every 20-30 minutes to mitigate fatigue. For the biofeedback group, trials alternated with and without the visual display in one-minute intervals. This approach aimed to maintain participants’ engagement in the biofeedback task, introduce moderate training variability, promote internalization of learned behaviors, and enable evaluation of immediate retention of biofeedback-induced gait changes.

After training, a post-test was conducted that repeated the six-minute pre-test trials in reverse order. All participants completed a Likert-scale questionnaire about exoskeleton use and training. Participants who received biofeedback also completed an abbreviated User Experience Questionnaire [34].

### Data and Statistical Analysis

Pre- and post-test outcomes were reported as mean values from the final three minutes of six-minute bouts. We use paired t-tests to assess differences in primary outcomes (metabolic cost and changes in targeted gait features) due to training within each group. Unpaired t-tests were applied for between-group comparisons during training and for pre-test versus post-test differences when both groups demonstrated a significant effect. Statistical parameter mapping was used to identify regions of significant training-induced change in ankle angle profiles.

To analyze adaptation, we performed regression analysis on metabolic cost, each of the four targeted gait features (step length, step width, ankle angle at 55% stride, and time of peak plantarflexion) and the variability of these gait features during assisted walking. For each outcome, we fit a dual-rate exponential of the form *y* = *c* + *a*_*fast*_*e*^*−t*/5^ + *a*_*slow*_*e*^*−t*/60^ to the minute-by-minute average or standard deviation of data during assisted walking. Fast and slow time constants of 5 and 60 minutes, respectively, were chosen based on prior adaptation studies [3,35]. The dispersion of each fitting parameter was estimated from non-parametric bootstrapping [36] and compared between groups using the Wilcoxon rank-sum test.

We additionally estimated total gait variability during the 60-minute training period. The standard deviation of step length and step width across all strides captured total variability related to foot placement. Total ankle angle variability was determined by calculating the standard deviation over all strides at each 0.1% of the gait cycle, then averaging across time points [37,38]. For participants who received biofeedback, we repeated these calculations for strides grouped across biofeedback-on trials and compared this to total variability during biofeedback-off trials using paired t-tests. We similarly compared energy consumption for trials when biofeedback was on versus off, using a first-order exponential fit to breath-by-breath data to estimate steady-state metabolic rate per trial [32].

Finally, to contextualize differences between observed and desired gait behaviors, we analyzed visual markers of error for the biofeedback task. The distance between real-time ankle angle and each target node was calculated in unitless form, with the x-axis scaled by stride duration and the y-axis scaled by the participant’s mean ankle range of motion during training. Step width error was calculated as a percentage of steps in which the arrow prompts were displayed. Adherence was assessed by comparing error markers in the final 30 seconds of each trial to those preceding biofeedback onset. Transient effects of biofeedback removal were evaluated by comparing the last 30 seconds of biofeedback trials to the first 15 seconds post-removal.

## Results

Training with biofeedback reduced the metabolic cost of walking with exoskeleton assistance (Fig. 2). The biofeedback group experienced a 23.5% ± 12.6% reduction in the metabolic cost of assisted walking due to training (paired t-test, p = 2e-5), while the control group showed a smaller, non-significant decrease of 11.8% ± 20.9% (paired t-test, p = 0.06). Inter-subject variability was substantial, particularly in the control group, where some participants experienced an increased metabolic cost after training (range: +1.2 to -0.94 W/kg). In contrast, all participants in the biofeedback group demonstrated energy savings (range: -0.13 to -1.92 W/kg).

**Figure 2.**
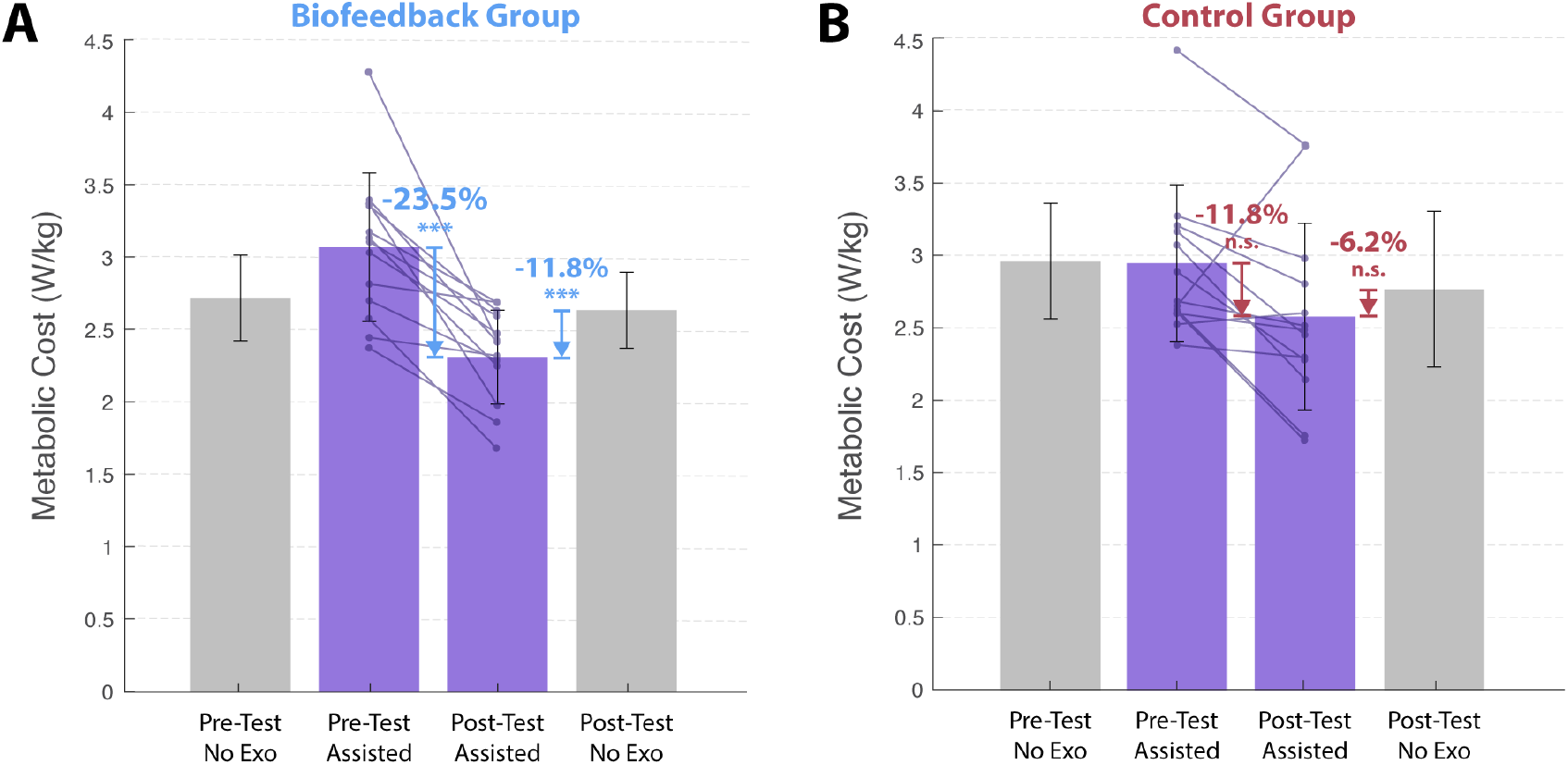
Effects of exoskeleton training on metabolic cost. Metabolic changes for **(A)** biofeedback and **(B)** control groups. Bars represent mean ±1 standard deviation for no-exoskeleton (gray) and assisted (purple) trials during pre- and post-tests. Individual subject data for assisted trials are overlaid.

The final metabolic benefit of walking with exoskeleton assistance was greater in the biofeedback group (Fig. 2). Given that part of the adaptation process entails learning to walk while wearing exoskeletons alone [5]—and this effect may have differed due to training—we compared assisted and no-exoskeleton trials for this analysis. Participants who received biofeedback training showed 11.8% ± 13.5% energy savings due to exoskeleton use (paired t-test, p = 0.01), compared to a non-significant reduction of 6.2% ± 19.4% in the control group (p = 0.3).

The characteristics of metabolic adaptation differed between groups. The evolution of metabolic cost over time was best described for the biofeedback group by *y* = 2.17 + 0.56*e*^*−t*/5^ + 0.65*e*^*−t*/60^, and for the control group by *y* = 2.53 + 0.87*e*^*−t*/5^ + 0.06*e*^*−t*/60^ (Fig. 3A). Each fitted parameter differed significantly between groups (Wilcoxon rank-sum test, p < 0.001; Fig. 3B). The larger contribution of the 60-minute component suggests earlier engagement of slower learning mechanisms for the biofeedback group compared to the control group. The estimated asymptote was also lower for the biofeedback group, indicating possible progression toward a lower steady-state metabolic cost.

**Figure 3.**
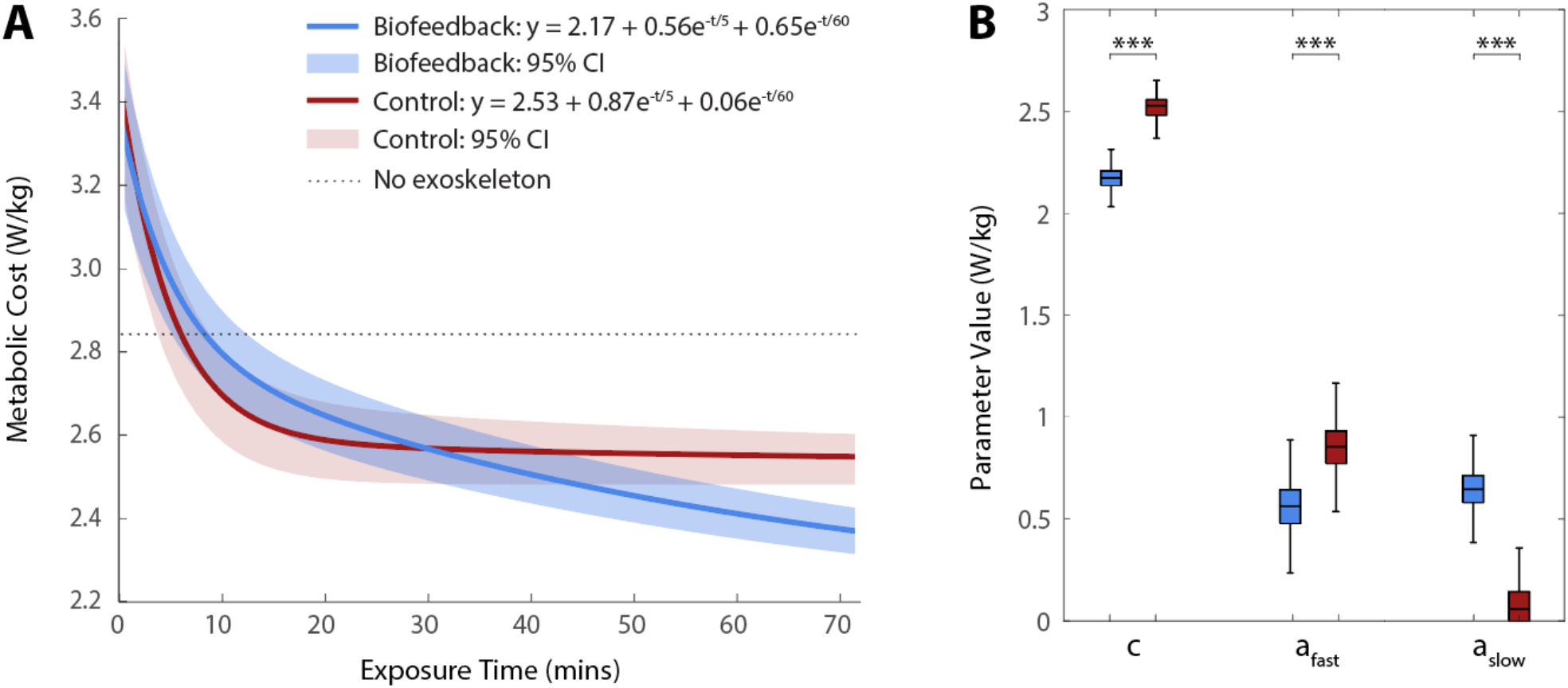
Metabolic adaptation. Dual-rate exponential (*y* = *c* + *a*_*fast*_*e*^*−t*/5^ + *a*_*slow*_*e*^*−t*/60^) fit to one-minute averages of breath-by-breath metabolic cost during assisted walking. **(A)** Exponential fits for biofeedback (blue) and control (red) groups, with 95% bootstrapped confidence intervals (CI). Dashed line shows nominal metabolic cost from the no-exoskeleton pre-test. **(B)** Interquartile range for each fitting parameter, with comparisons between groups using Wilcoxon rank-sum tests (*** for p < 0.001).

Inclusion of biofeedback increased the effectiveness of training at eliciting the targeted gait modifications (Fig. 4). Participants who trained with biofeedback significantly increased step length (+4.6 ± 7.1 cm; paired t-test, p = 0.01), while the control group showed no significant change (+0.6 ± 4.4 cm, p = 0.7). The effect of training on step width was not statistically significant for either group. However, adjustments in the desired direction appeared larger (Fig. 4C) and more rapid (Fig. 4D) for the biofeedback group (−2.0 ± 1.8 cm, p = 0.2) compared to the control group (−1.1 ± 2.2 cm, p = 0.4). Ankle dorsiflexion preceding push-off significantly increased for the biofeedback group (4.7 ± 4.1°, p = 0.04), and did not change significantly for the control group (3.0 ± 2.8°, p = 0.07). Mid-stride ankle angle continued to change more rapidly in the biofeedback group, suggesting faster convergence toward optimal behavior (Fig. 4F). Neither group demonstrated a reliable effect of training on peak plantarflexion timing (biofeedback: +1.1 ± 2.0% stride, p = 0.02; control: +1.0 ± 0.9% stride, p = 0.1). Differences due to training were statistically significant for the biofeedback group, but high inter-subject variability and comparable means between groups decrease confidence in this effect. Interpretation of these timing changes are further complicated by differences in stride duration varying between groups (Fig. 4A)

**Figure 4.**
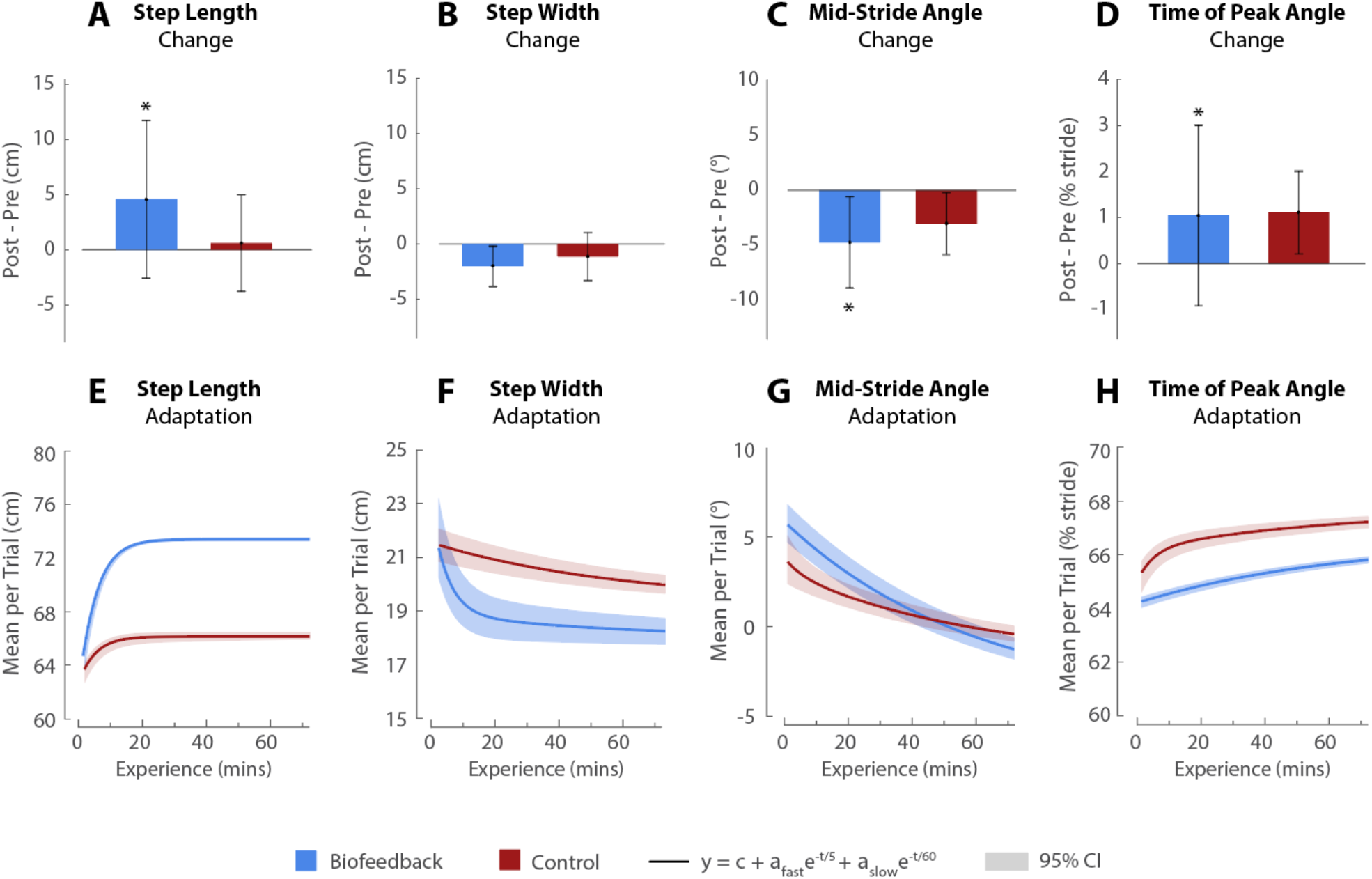
Gait adaptations. Changes in targeted gait features during assisted walking for the biofeedback (blue) and control group (red). **(A-D)** Bars indicate mean ± 1 standard deviation of gait changes between pre-test and post-test. **(E-H)** Curves represent dual-rate exponential models fit to the mean value of each gait feature during one-minute trials. Shaded regions indicate 95% bootstrapped confidence intervals. Model fits for biofeedback and control groups, respectively: (E) Step length: *y* = 1/(0.014 + 0.0022*e*^*−t*/5^), *y* = 1/(0.015 + 0.00073*e*^*−t*/5^). (F) Step width: *y* = 17.97 + 3.19*e*^*−t*/5^ + 0.93*e*^*−t*/60^, *y* = 19.31 + 2.19*e*^*−t*/60^. (G) Mid-stride ankle angle: *y* = −4.36 + 10.26*e*^*−t*/60^, *y* = −1.93 + 0.81*e*^*−t*/5^ + 5.03*e*^*−t*/60^. (H) Time of peak angle: *y* = 1/(0.015 + 0.00054*e*^*−t*/60^), *y* = 1/(0.015 + 0.00029*e*^*−t*/5^ + 0.00033*e*^*−t*/60^). Fitted parameters differed between groups for all features (Wilcoxon rank-sum tests, p < 0.001).

The selected ankle angle features reflected broader changes in ankle joint kinematics around push-off. Both types of training significantly altered ankle angle trajectories: from 49.0% to 59.6% of stride for the biofeedback group (SPM, p = 0.003) and 52.2% to 60.4% of stride for the control group (p = 0.008). No regions of significant difference were observed for between-group comparisons of ankle angle trajectories. However, within-group comparisons indicate that participants who trained with biofeedback-initiated changes in ankle behavior earlier and sustained them over a longer portion of the gait cycle.

Participants responded quickly to biofeedback cues, but reductions in biofeedback task error did not consistently yield intended gait changes. During participants’ first one-minute exposure to biofeedback, step width error decreased from 53% to 17% (paired t-test, p = 0.003), then partially rebounded to 34% post-removal (p = 0.02). However, the average step width was 2 cm narrower than the target value during active biofeedback, suggesting that participants varied between overly narrow and wide steps rather than achieving a stable, optimal step width. In addition, total ankle angle error decreased from 14% to 10% during initial biofeedback exposure (p = 0.08) and remained at 10% after biofeedback removal (p = 0.88). However, this error reduction was not due to closer matching of ankle angle profiles to intended angle features. In fact, biofeedback exposure resulted in a mid-stride ankle angle that was 1.4° more plantarflexed and a peak angle that occurred 1% earlier in the gait cycle compared to walking without biofeedback. Instead, participants appeared to reduce visual error through a compensatory strategy of increasing step length by 4.0 cm, exceeding the targeted value by 3.2 cm.

Gait variability during the 60-minute training period was generally higher for the biofeedback group than the control group. When training included biofeedback, step length variability was significantly greater (4.0 ± 0.9 cm vs. 2.5 ± 0.7 cm; unpaired t-test, p = 9e-5), as was ankle angle variability (4.7 ± 1.1° vs. 1.1 ± 0.8°, p = 0.02), though step width variability was comparable between groups (biofeedback: 2.4 ± 0.6 cm, control: 2.2 ± 0.7 cm, p = 0.5). For the biofeedback group, elevated gait variability persisted even during trials when biofeedback was removed. Ankle angle variability was similar between trial types (active: 4.8 ± 1.1°, inactive: 4.4 ± 1.2°; paired t-test, p = 0.11), as was step width variability (active: 3.7 ± 0.9 cm, inactive: 3.8 ± 1.0 cm, p = 0.2). Step length variability decreased after biofeedback removal, but remained elevated (active: 5.0 ± 1.0 cm, inactive: 4.3 ± 0.8 cm, p = 1e-6).

Regression analysis further indicates groupwise differences in exploratory gait behaviors. Fit parameters for gait variability analyses differed significantly between groups for all measures (Wilcoxon rank-sum test, p < 0.001). The biofeedback group generally exhibited greater exploration and change toward stable behaviors, while the control group showed lower and less dynamic variability. This pattern was observed for variability in step length (biofeedback: *y* = 2.32 + 0.89*e*^*−t*/60^, control: *y* = 1.82 + 0.80*e*^*−t*/5^ + 0.48*e*^*−t*/60^), step width (biofeedback: *y* = 1.74 + 0.27*e*^*−t*/60^, control: *y* = 1.63), and mid-stride ankle angle (biofeedback: *y* = 2.99 + 1.29*e*^*−t*/60^, control: *y* = 2.52 + 0.19*e*^*−t*/5^ + 0.49*e*^*−t*/60^). This pattern was not evident for variability of peak plantarflexion timing (biofeedback: *y* = 3.0 + 1.3*e*^*−t*/60^, control: *y* = 2.5 + 0.2*e*^*−t*/5^ + 0.5*e*^*−t*/60^), though overlapping confidence intervals suggest comparable adaptation.

Perceptions of exoskeleton use were modest in both groups, while the biofeedback display was viewed favorably by its users. Exoskeleton usability ratings were moderate, with large improvements reported by the end of the session due to experience (Table 1). These ratings were similar across groups, despite differences in training protocols. The biofeedback display was positively received, with a majority of participants rating it as motivating, exciting, enjoyable, understandable, and supportive (Fig. 5). There were mixed perspectives on how easy the biofeedback task was to learn, with ratings skewed positive on average (Fig. 5D).

**Table 1.** Questionnaire responses on exoskeleton use. Impressions of walking with exoskeletons (0: strongly disagree, 1: disagree, 2: neither agree nor disagree, 3: agree, 4: strongly agree) and perceived changes from the beginning to the end of the session (0: decreased significantly, 1: decreased, 2: no change, 3: increased, 4: increased significantly).

| Survey Question | Biofeedback | Control |
| --- | --- | --- |
| It felt easier to walk when the device was active. | $2.8 \pm 1.0$ | $2.7 \pm 0.8$ |
| I felt in control of my movements when the device was active. | $2.3 \pm 0.9$ | $2.1 \pm 0.9$ |
| I feel comfortable using the device. | $2.5 \pm 1.0$ | $2.5 \pm 0.9$ |
| I thought the device was easy to use. | $2.4 \pm 1.0$ | $2.6 \pm 1.0$ |
| I think that I would like to use this device frequently. | $1.3 \pm 1.3$ | $1.3 \pm 1.3$ |
| I found the device unnecessarily complex. | $1.2 \pm 0.7$ | $1.1 \pm 0.6$ |
| I felt very confident using the system. | $2.6 \pm 0.7$ | $2.4 \pm 0.7$ |
| Change in how easy it felt to walk with the device. | $3.6 \pm 0.5$ | $3.7 \pm 0.5$ |
| Change in comfort with the device. | $3.4 \pm 0.5$ | $3.1 \pm 0.9$ |
| Change in feeling of control over movements with the device. | $3.1 \pm 0.5$ | $3.1 \pm 0.6$ |

**Figure 5.**
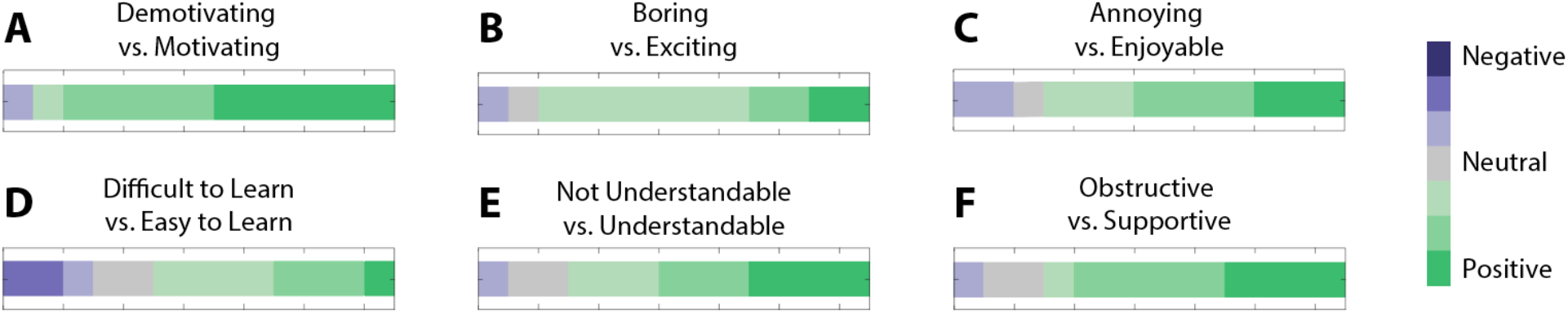
Questionnaire responses on biofeedback design. Participants selected between descriptive adjectives and their antonyms to assess the visual display’s **(A-B)** stimulation, **(C)** attractiveness, **(D-E)** perspicuity, and **(F)** dependability [33]. Stacked bar charts are shown of 1-7 Likert scale responses, with 1 (dark purple) as most negative, 4 (gray) as neutral, and 7 (dark green) as most positive. Intermediate values are represented by intermediate colors.

## Discussion

Biofeedback was highly effective at facilitating motor learning with ankle exoskeletons. The use of this approach made one hour of training twice as effective, resulting in 12% (0.42 W/kg) lower energy consumption while walking with exoskeletons compared to walking in normal shoes (Fig. 2). These savings are functionally meaningful—equivalent to removing a 9 kg backpack while walking [39]—and are comparable to expert-level performance achieved in prior multi-day studies [3]. The minimum metabolic cost observed during assisted walking, as well as the energy savings relative to no-exoskeleton and zero-torque conditions, were previously reported after four to five days of training [3]. These results indicate that biofeedback can provide about a fourfold reduction in the time required to attain expert-level energy savings with an ankle exoskeleton. Our analysis also suggests that the biofeedback group was progressing toward a lower steady-state metabolic cost than has been previously reported (Fig. 3) [3]. Longer data collection is needed to definitively determine whether this biofeedback approach can unlock greater benefits from exoskeletons, and if so, to what extent. By helping users benefit more quickly—and potentially also to a greater degree—this approach demonstrates the potential value of biofeedback-based training as a tool to enhance mobility.

The utility of biofeedback may depend on the intended duration of exoskeleton use. Biofeedback incurred an initial metabolic penalty, potentially due to increased cognitive demand and gait variability [40,41]. However, this was offset after approximately 30 minutes of use as individuals discovered more efficient gait strategies (Fig. 3). Participants who trained with biofeedback continued to significantly decrease their metabolic cost, rather than plateau within 30 minutes of use, reflecting that slow learning processes, such as improved sensory prediction, were engaged earlier and progressed further in the biofeedback group [42]. Given that our study modestly exceeded the time constant of the modeled slow phase, longer-duration studies would help validate and refine the trends observed here.

Biofeedback appeared to facilitate energy improvements by promoting beneficial gait changes. Participants who were exposed to biofeedback demonstrated more gait exploration with respect to step length and ankle joint kinematics. This behavior persisted even when biofeedback was removed, suggesting that participants were developing intrinsic models of learned behavior. Gait exploration led to equivalent or greater changes in the four targeted features: lengthening and narrowing steps while dorsiflexing more preceding a later push-off (Fig. 4A-D). In most cases, the rate of gait adaptation was faster for the biofeedback group and more closely approached expected energy-optimal characteristics (Fig. 4E-H). However, participants who trained with biofeedback did not yet converge on the gait patterns observed in fully adapted users [5]. Gait variability also remained elevated and non-steady-state, indicating that the biofeedback group had not yet acquired stable motor skill.

Anecdotal evidence suggests that biofeedback may be particularly effective at correcting maladaptive responses to new motor contexts. Participants for whom assisted walking was initially most taxing benefited most from biofeedback, while in the control group, participants with the highest initial costs followed the group average (Fig. 2). Separately, we observed an illustrative case in which pair of participants—one from either group—demonstrated the potential of biofeedback to mitigate the maladaptive response of taking progressively shorter steps in a marching gait. Both subjects began with an exoskeleton-assisted step length of approximately 63 cm and were expected to adapt to a 70 cm step length. The subject who trained with biofeedback explored step lengths up to and beyond 70 cm and achieved a final energy reduction of 0.76 W/kg (23%). In contrast, the control subject never exceeded a step length of 66 cm and further shortened their steps to 57 cm, with energy cost increasing by 1.16 W/kg (44%) in the post-test. From these cases, we posit that if the demands of biofeedback are sufficiently high to warrant sparing use, the most strategic approach would be to deploy biofeedback as an intervention when maladaptive behaviors are detected.

While effective at promoting desirable metabolic and kinematic changes over the session, our biofeedback design was limited in its ability to acutely induce desired outcomes. Gait characteristics were not consistently at target values and metabolic cost was not consistently lower during trials with active biofeedback. The latter may be attributed to increased cognitive load, suboptimal combinations of targeted gait features, and/or non-economical compensations above the ankle. It is possible that consistently inducing low-energy-cost gaits would have further improved the rate and extent of motor learning. Nevertheless, our findings suggest that biofeedback can effectively promote beneficial changes even without direct exposure to optimal behaviors. Future research on motor learning for gait should focus on identifying and guiding exploration in task-relevant dimensions, even when the underlying cost landscape is difficult to define.

Several design elements may explain discrepancies between intended and observed gait behaviors. First, step width feedback lacked sufficient resolution to support fine adjustments, leading to oscillations between too-narrow and too-wide steps. Second, combining visualizations of stride duration and ankle angle obscured changes in both measures. Participants appeared to focus on lengthening their steps to reduce visual error, likely because this dimension was easier to understand and modulate, which mitigated beneficial changes in ankle kinematics. These findings underscore the need for high-specificity feedback. If multiple task-relevant dimensions are to be combined, step length and width may be better represented by a visualization of actual versus desired foot placement. Additionally, although biofeedback targets were selected from prior expert behavior [5], these were based on average trends and may not have been ideal for all participants or for early-stage training. Future approaches could involve adaptive targets, shifting over time as users improve [20]. These refinements could enhance the ability of biofeedback to elicit desired gait changes.

Several experimental limitations also affected the generalizability of our findings. Although this study was well-powered to detect training-related changes, its unpaired design and high inter-subject variability prevented statistical analysis with respect to initial physiological or biomechanical differences. We also did not systematically assess the effectiveness of our feedback delivery strategy, including the impact of frequent one-minute trials and the limited nature of a three-minute washout preceding final measurements. We were further unable to systematically isolate the mechanisms underlying the observed metabolic changes. Finally, because data were collected in a single session, additional research is needed to assess the retention of these training effects. Promisingly, prior research suggests that accelerating the slow phase of learning promotes faster re-learning after periods of nonuse [43,44], highlighting another potential advantage of the biofeedback-induced changes we observed. Addressing these points will help clarify the practical relevance of our approach for long-term exoskeleton use across a wider range of individuals.

Translating our findings to real-world applications presents several design challenges. One key consideration is the balance between information density and cognitive demand. We prioritized providing detailed and comprehensive information, and while healthy young adults found our biofeedback system to be motivating and supportive, they had mixed perspectives on how easy it was to learn (Fig. 5). A potential alternative strategy to reduce concurrent demands without loss of information would be to vary feedback to target one goal behavior at a time (e.g., prioritizing the gait dimension with the largest deviation from desired outcomes). This approach may improve skill acquisition with lower cognitive demand within each mode. However, frequent mode switching could increase cognitive load and impair performance overall [45]. Formalizing the trade-offs of such approaches would inform next steps for the design of multi-dimensional biofeedback paradigms. These insights will be critical for the extension of this approach to individuals with slower motor learning, such as those neurological impairments or age-related declines. Additionally, to support real-world use, future research should investigate modalities such as auditory or haptic cues and how they can be made most effective given high task complexity [21]. Many open questions remain regarding how to develop effective and tractable training approaches with biofeedback for these complex use cases.

Finally, this study highlights the need for interventions that not only improve device effectiveness, but also positively impact users’ attitudes towards assistive devices. Participants reported similar perceptions of usability, ease of use, and comfort with the exoskeletons, regardless of training type or measured changes in energy economy. Therefore, despite the energetic benefits conferred by this intervention, it is unlikely to affect device desirability or adoption on its own. This may differ for individuals with greater difficulty walking without assistance. Understanding what metabolic effects are detectable and meaningful for people of different ages, fitness and mobility levels could critically guide assistive device adoption.

## Conclusions

Targeted biofeedback can significantly accelerate learning how to walk with an exoskeleton. This study demonstrates that biofeedback is effective not only in structured tasks with explicit goals, but also in complex motor contexts where adaptation is typically slow, the potential action space is large, and the underlying cost landscape is difficult to characterize. While feedback may be most effective when it guides individuals directly toward optimal behaviors, meaningful improvements can still be achieved by promoting changes along task-relevant dimensions, particularly when the direction of improvement can be identified. By guiding novice exoskeleton users to modify their gait kinematics in directions previously associated with reduced metabolic cost, we enabled rapid improvements in exoskeleton-assisted walking. These gains, which were comparable to expert-level energy savings, suggest that biofeedback can expedite the slower phase of motor learning and enable users to benefit from assistive devices more quickly than with conventional training alone. Our findings position biofeedback as versatile strategy to enhance learning in less-structured motor contexts, paving the way for faster gait rehabilitation, more intuitive human-robot interaction, and expanded use of mobility-enhancing technologies.

## Acknowledgments

This work was supported by the National Science Foundation (1828993 and DGE-1656518) and the Gabilan Stanford Graduate Fellowship.

